# Evidence for the presence and diagnostic utility of SPM in human peripheral blood

**DOI:** 10.1101/2022.04.28.489064

**Authors:** Jesmond Dalli, Esteban A Gomez, Charles N Serhan

**Affiliations:** William Harvey Research Institute, Barts and The London School of Medicine and Dentistry, Queen Mary University of London, Charterhouse Square, London, EC1M 6BQ UK; Centre for Inflammation and Therapeutic Innovation, Queen Mary University of London, London, UK; Center for Experimental Therapeutics and Reperfusion Injury, Department of Anesthesiology, Perioperative and Pain Medicine, Brigham and Women’s Hospital and Harvard Medical School, Boston, Massachusetts 02115, USA

## Abstract

We thank O’Donnell *et al*, for their comments on our contribution and are grateful to be afforded this opportunity to formally respond to their critique^24^.

We are surprised by the author’s assertion relating to the biological relevance of SPM because a simple literature search for related terms such as ‘resolvin’ in PubMed yields an abundance (>1,420 publications) of evidence supporting the potent biological activities and the diagnostic potential of some of these mediators. Several co-authors of the O’Donnell’s *et al* manuscript, have published on the resolvins and SPMs, including some publications within recent weeks. Importantly, O’Donnell *et al*, misreport as well as mis-apply criteria for peak identification reported in the Gomez *et al*, publication which lead to the flawed analysis they performed.

In this response therefore, we provide a step-by-step clarification of the methodologies used in Gomez *et al*, and a side-by-side comparison of the underlying data to clarify any confusion. We also demonstrate that using the orthogonal criteria discussed by O’Donnell *et al*, we obtain essentially identical results thus providing additional validation of our techniques and support the conclusions.

## Main Text

O’Donnell *et al*,assert that because they can achieve an area under the curve (AUC) of ≥ 2000 cps by integrating background noise, our criteria ‘*lead to flawed methodology to SPM analysis brings into question the very occurrence of many of these lipids in biological samples, their proposed impact on inflammatory processes, and biomarker claims’*.

This assertion is based on the misapplication of the criteria that were describe clearly, in our view, in the methods section of Gomez *et al*, leading to erroneous results and therefore conclusions. In Gomez *et al*, the criterion described for peak identification and integration and therefore calculation of area under the curve (AUC) requires the presence of a distinct peak in the chromatogram as denoted by the following text in the methods: *presence of a* ***<u>peak</u>*** *with a minimum area of 2000 counts*. In the chromatogram provided by O’Donnell *et al*, there is <u>no discernible peak</u> and therefore it would not meet the basic criterion for integration. To further clarify the application of the criteria employed in the analysis of data presented in Gomez *et al*, we provide an illustration that presents the decision pathway we use for peak identification and integration (Supplemental Figure 1). We also provide examples of chromatograms from our data analysis which report peaks that were used for the calculation of concentrations of distinct mediators. For these examples we selected those samples where the AUC of the peak of interest was close to the 2000 cps cut-off as a further illustration of minimum signals employed for identification and quantitation of the lipid mediators reported in this publication (Supplemental Figure 2).

O’Donnell *et al*, also argue that because we did not use signal-to-noise (s/n) ratios as cut-off parameters for determining the lower limits of quantitation (LLOQ) our analysis is flawed and further that SPMs do not therefore exist in biological systems. We respectfully disagree with this assertion and point of view. Firstly, because as detailed above, there is extensive documentation from many groups (including some of the co-authors of *O’Donnell et al* ^1, 2, 3, 4, 5, 6, 7, 8^) which identify and quantitate SPM in an array of biological systems. Indeed, as discussed above, a PubMed search term of “resolvins” gives 1,420 reports (as of today) from groups around the world using commercially available resolvins and other SPMs confirming their potent stereoselective actions in a wide range of models and presence in human tissue samples ^3, 9, 10, 11, 12, 13, 14, 15, 16^. Secondly, while it is the case that the different entities mentioned by the authors have recommended the used of s/n ratios as an analytical criterion, the suggestion made by O’Donnell *et al*,that this is the only criterion recommended by these entities is not fact. Indeed, these entities identify a number of different approached for the quantitation of molecules. Furthermore, several of these entities even state that these methods have their limitations and that they may not be applicable to all target molecules. What *is* universally accepted is the need for the signal (peak) of interest to be sufficiently different from the background to allow for the robust and reproducible quantitation of the molecule.

Whilst we acknowledge that using s/n ratios can be useful in identifying the LLOQ for a particular assay, their application is not universally accepted and has a number of limitations. Firstly, the calculation of this parameter is influenced by instrument configuration (e.g. the type of detector) and data processing approaches (https://sciex.com/content/dam/SCIEX/pdf/tech-notes/all/mass-spectrometry-cms_059150.pdf). Furthermore, a review of the literature, including documents cited by O’Donnell *et al*, demonstrates that there are several approaches for the calculation of such parameters which may lead to different outcomes, and there are also different guidelines for cut-offs to be employed^17, 18^. Indeed, as the authors themselves state, there is still no universally accepted consensus in their field on the cut-off value, and the LLOQ of 5 was set arbitrarily, with other investigators adopting a lower cut-off value (*e*.*g*., s/n 3) whilst others opted for a higher cut off value (*e*.*g*., s/n 10).

A further complicating factor that all workers in this field are acutely aware of is represented by the presence of several isobaric isomers which elute in the proximity of the mediators of interest. This aspect can be appreciated in many of the chromatograms provided in Supplemental Figure 2. These isomers interfere with the calculation of s/n ratios since their presence precludes the identification of an appropriate region within the chromatogram that represents a true baseline, and which is therefore sufficiently wide accurately to calculate the value for the ‘noise’.

To overcome this limitation, we developed a method that retains the required robustness and is not hampered by the presence of biological isomers within the chromatogram. In support of the robustness of our method we provided tables in our publication where we present the coefficient of variations on the two instruments we employed for the analysis (Supplemental Tables 11 and 12 from Gomez *et al*, publication). Unfortunately, it appears that O’Donnell *et al*, have overlooked this key dataset. Taken together, this evidence contradicts the assertions made by O’Donnell *et al*.

In the interests of extending this scientific discussion as well as demonstrating the robustness of our approach, we have reanalysed the underlying data from the Gomez *et al* publication, this time applying an LLOQ with a s/n ≥ 5. Due to 1) space limitations 2) since the data underlying this Figure 1 in the Gomez *et al* publication represents the vast majority of the data presented in Gomez *et al*, 3) given that O’Donnell *et al*, claim that our approach does not support the utility of measuring SPMs as biomarkers and 4) since machine learning models are exquisitely sensitive to changes in the variables being used, in this response we report the results obtained from the reanalysis of data presented in Figure 1 of Gomez *et al*. For this analysis we employed the *relative noise concept*, which calculates the baseline under the peak of interest by calculating the noise in the chromatographic region. This concept overcomes many of the issues classically associated with s/n calculations, including the subjective nature of the noise region selection and the presence of other peaks within the region of interest that make it challenging to identify a sufficiently wide ‘noise’ region to obtain a correct estimate. The following documentation details this concept and the underlying algorithm (<u>SCIEX OS for Triple Quadrupole Systems Software User Guide</u>).

**Figure 1:**
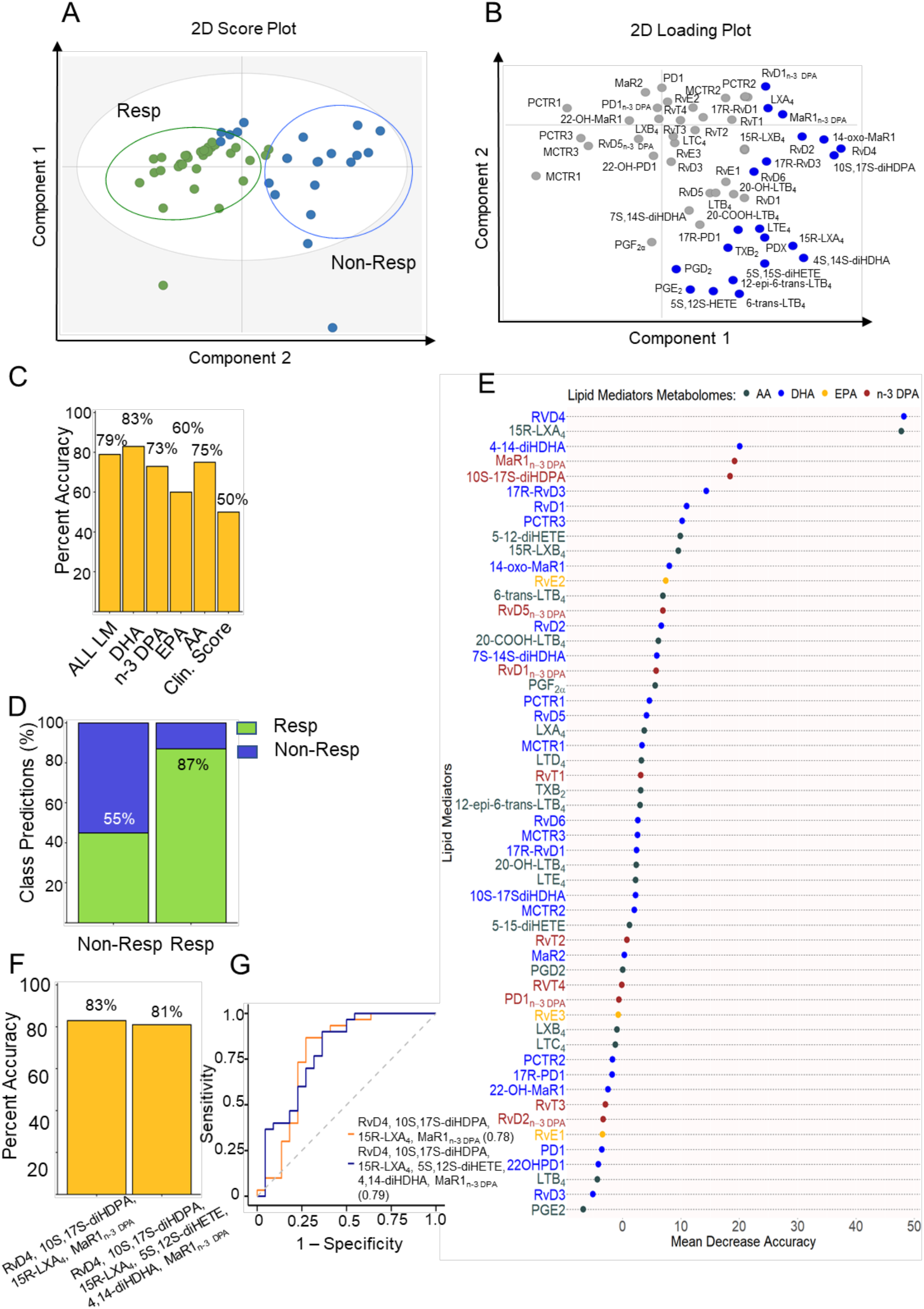
Reanalysis of baseline plasma lipid mediator profiles supports their potential utility as biomarkers. Plasma was collected from RA patients prior to the initiation of treatment with DMARDs and lipid mediator concentrations established using LC-MS/MS based lipid mediator profiling (see methods for details). (A,B) OPLS-DA analysis of peripheral blood lipid mediator concentrations for DMARD responders (Resp) and DMARD non-Responders (Non-Resp). (A) 2-dimensional score plot with the gray circle representing the 95% confidence regions. (B) 2-dimensional loading plots. Lipid mediators with VIP score greater than 1 are highlighted in blue and upregulated in Non-Resp. Results are representative of n = 30 Resp and n = 22 Non-Resp. (C) % accuracy score of prediction models based on the combination of all lipid mediators identified and quantified (AL LM) or individual fatty acid metabolomes as indicated. Clin. Score = clinical score (see methods from Gomez *et al*, for parameters included). (D) Classification predictions for each class (sensitivity and specificity) of the n-3 DPA model. Green indicates the samples that were predicted as Resp while blue indicates those patients predicted Non-Resp. Percentages indicate true positives (Resp class) and true negatives (Non-Resp class). (E). Relevance of lipid mediators in the prediction performance of the “ALL LM” model based on decreasing accuracy. (F) % accuracy score of models using the indicated SPM. (G) Receiver operating characteristic (ROC) curves and AUC values for predictive models based on the indicated SPM. All the models were created using the random forest methodology (“randomForest” package from R).

As seen from data presented in Supplemental Figure 2, side by side comparison of s/n ratios and our criteria demonstrates that these fulfil the minimum s/n criteria of ≥ 5. Indeed, these results demonstrate that peaks with a 2000 AUC value easily achieve a s/n value that is > 5. Assessment of lipid mediator concentrations obtained using this orthogonal approach using partial least square discriminant analysis demonstrated a separation between the clusters representing DMARD responsive and unresponsive patients. Furthermore, evaluation of potential lipid mediator biomarkers using machine learning identified essentially the same lipid mediators as the potential predictors for treatment outcome. This result was further corroborated by evaluating the concentrations of these molecules in the second cohort of patients, analysis that once again was performed using the s/n ≥ 5 cut-off. Of note both the accuracy and AUC values obtained in this reanalysis of the validation cohort gave essentially identical outcomes supporting the robustness of the analysis preformed in Gomez *et al*. (Figure 1).

O’Donnell *et al*. also asserted that the MS/MS criteria using in Gomez *et al*. employed in the identification of SPMs were flawed and that the ions employed in the identification were derived from background noise. They provide MS/MS spectra that were generated in their laboratory using different instrumentation, and likely different instrument parameters, and then proceed to use these as reference spectra for their argument. They further used MS/MS spectra from a reference database which was compiled using completely different mass spectrometers, and different acquisition parameters including different cut-off values for the MS/MS spectrum (which go below an m/z of 100), to further bolster their argument.

We are surprised that the authors deem this to be an appropriate approach given that they themselves acknowledge in their article that different mass spectrometers are likely to yield distinct MS/MS spectra. We therefore find this argument to be flawed. O’Donnell *et al*, then go on to claim that they were able to obtain a spectrum that would match that of Maresin (MaR) 1 from a blank sample. It is obviously impossible to guess what contaminants there may have been in their instrumentation that may have contributed to their result or where within the chromatogram they obtained the spectrum displayed in the figure (i.e., was it within the region where this mediator would be expected to elute?). Furthermore, as can be observed from data presented in Supplemental Figure 3, evaluation of blanks on our instrumentation did not yield MS/MS spectra that contained ions that could be linked with lipid mediator identification. Focusing on MaR1, the example provided by O’Donnell *et al*, when we extracted ions with an m/z of 359.4 corresponding to MaR1 parent ion (and other dihydroxylated SPM derived from DHA) we did not observe any eluting at the retention time corresponding with that of MaR1 (and other dihydroxylated SPM). Furthermore, evaluation of MS/MS spectra captured for molecules eluting before and after the retention time of MaR1 did not yield any of the ions reported by O’Donnell *et al*. suggesting that the spectra they reported arise from contaminants within their instrumentation.

Whilst we concur with O’Donnell *et al*. that robust criteria for the identification of SPM are required we believe that the description they provide of a ‘*generally similar pattern of relative abundance’*, for MS/MS spectral matching is subjective at best and does not fulfil the rigorous standards that they suggest are important.

We agree with O’Donnell *et al*, that it is not possible to obtain MS/MS spectra that are of diagnostic value from every sample for every analyte. This due to several factors primarily relating to the data acquisition process in cases where in LC-MS/MS experimental data is acquired ‘on the fly’. Since many of the analytes elute close to each other, the instrument dwell times make acquisition of data for all analytes challenging.

We disagree with their assertion that “*when lipids are present in very low amounts in a biological sample, MS/MS spectra are often inconclusive and do not compare well with standards”*. Firstly, because the use of rigorous MS/MS spectra and deduction of these very diagnostic ions together with evaluation of their biological activities are central in the structure elucidation of bioactive mediators. Importantly the identity of these molecules was confirmed using total organic synthesis^19^ and their presence and potent bioactions confirmed by a large number of groups in a diverse array of biological systems, including some of O’Donnell’s co-authors^1, 2, 3, 4, 5, 6, 7, 20, 21, 22^. Secondly, we consider that MS/MS spectra derived from a subset of the samples provides additional confirmation for the presence of the analytes of interest.

To amplify and support this point in the interest of scientific discussion, we have reanalysed the underlying the data presented in Gomez *et al*. employing the library match function in Sciex OS, whereby the software matches the ions present in the samples with those in the reference spectrum and provides a score. Here we used a score of ≥70% and matching retention time as a confirmation of a positive match. As can be appreciated in the examples provided in Supplemental Figure 4, this reanalysis confirmed the initial observations published in Gomez *et al*. that many of the SPMs are present in plasma from RA patients.

## Summation

Whilst we welcome genuine scientific discussion and critical evaluation of published work, it is clear that the assertions made by O’Donnell *et al*. are not shared by the many investigators (https://pubmed.ncbi.nlm.nih.gov/?term=resolvins&sort=date). Moreover, we find these assertions unfounded, erroneous and flawed because: 1) their argument is based on the misapplication of a key criterion described in Gomez *et al*. and 2) a reanalysis using the orthogonal criteria, namely those described by O’Donnell *et al*. yielded essentially the same results and support the conclusions published in Gomez *et al*. This further substantiates both the presence of SPMs in human peripheral blood, in line with findings made by others ^1, 3, 4, 16, 21, 23^, and their utility as biomarkers. The value of peer review cannot be under appreciated and we’ve welcomed the opportunity to discuss results published in Gomez *et al*. and further demonstrate the strength of the identifications and conclusions regarding the SPMs and other lipid mediators e.g., eicosanoids including LTB_4_, and certain prostaglandins in this study.

## Methods

Data acquisition, multivariate analysis and machine learning models were performed as detailed in Gomez *et al*. For the calculation of signal-to-noise ratios the AutoPeak and Noise filtering, and Relative Noise functions in Sciex OS v2.1 were employed.

## Supplementary Figures

**Supplementary Figure 1: Step by step illustration of the criteria for peak integration**.

**Supplementary Figure 2: Representative examples of peaks corresponding to distinct mediators identified in Gomez *et al***. In these examples we also denote the AUC values and the corresponding sign-to-noise ratios. Furthermore, one can also appreciate the presence of several biological isomers that elute in close proximity to many of the peaks of interest.

**Supplementary Figure 3: Blank samples do not yield MS/MS spectra of diagnostic value**. (A) Screenshot of MaR1 standard denoting the retention time of this mediator (B-D) A blank was injected and the (B) extracted total ion chromatogram for m/z 359.4 corresponding to the MaR1 parent ion was obtained. (C) MS/MS spectrum obtained from the signal reported at a retention time of 13.93 in the ion chromatogram reported in (B) demonstrating the absence of diagnostic ions.

**Supplementary Figure 4: Matching of MS/MS spectra from obtained from RA patient plasma samples using the library function in Sciex-OS confirms the presence of these autacoids**. MS/MS spectra obtained from plasma samples described in Gomez *et al*, were matched using the library function in Sciex-OS and a threshold value >70% score.

