## Supplemental figures for "Evidence for the presence and diagnostic utility of SPM in human peripheral blood"

Supplemental Figure 1

Is there a distinguishable peak?

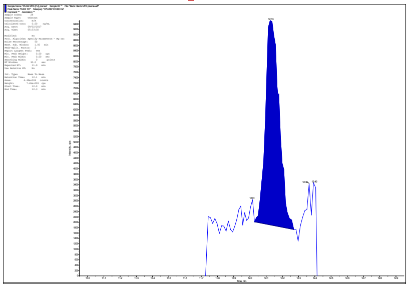

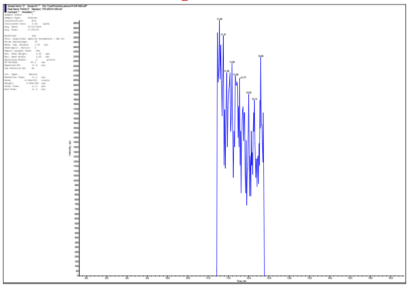

YES → Does the peak elute at the same retention time as that of the relevant standard ?

NO → No integration

Does the peak elute at the same retention time as that of the relevant standard ?

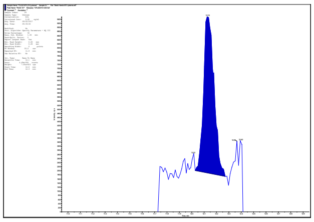

YES → Is the area ≥ 2000 counts?

NO → No integration

Is the area ≥ 2000 counts?

YES → Quantitation

NO → **No** quantitation

### Supplemental Figure 2

#### RvD2

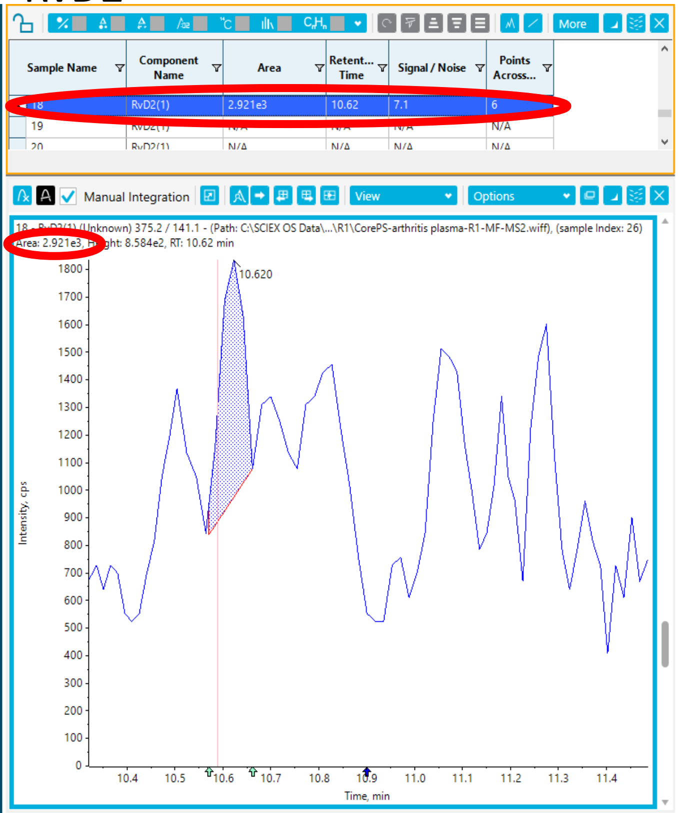


#### RvD3

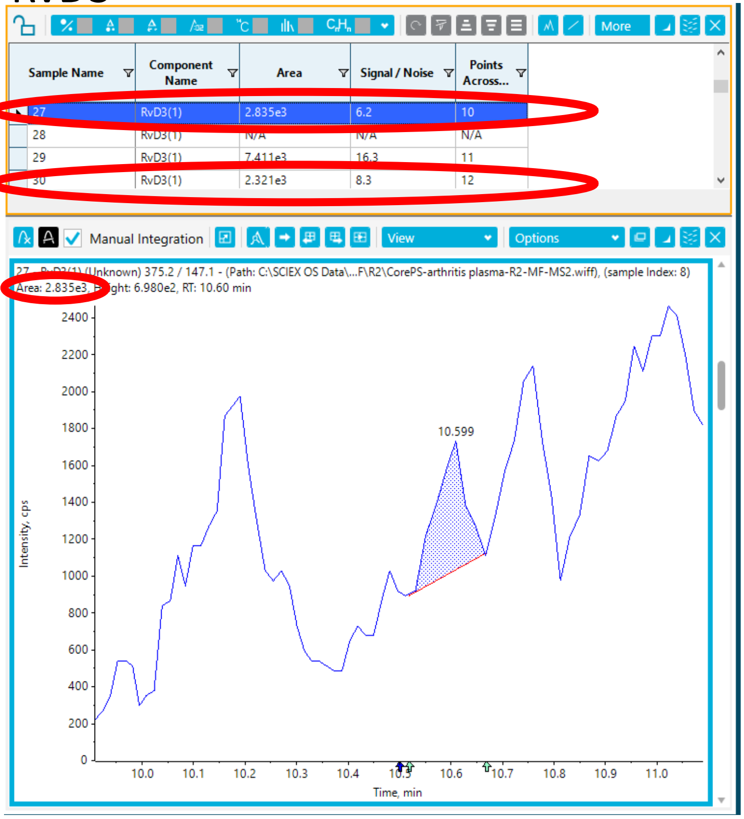


#### RvD4

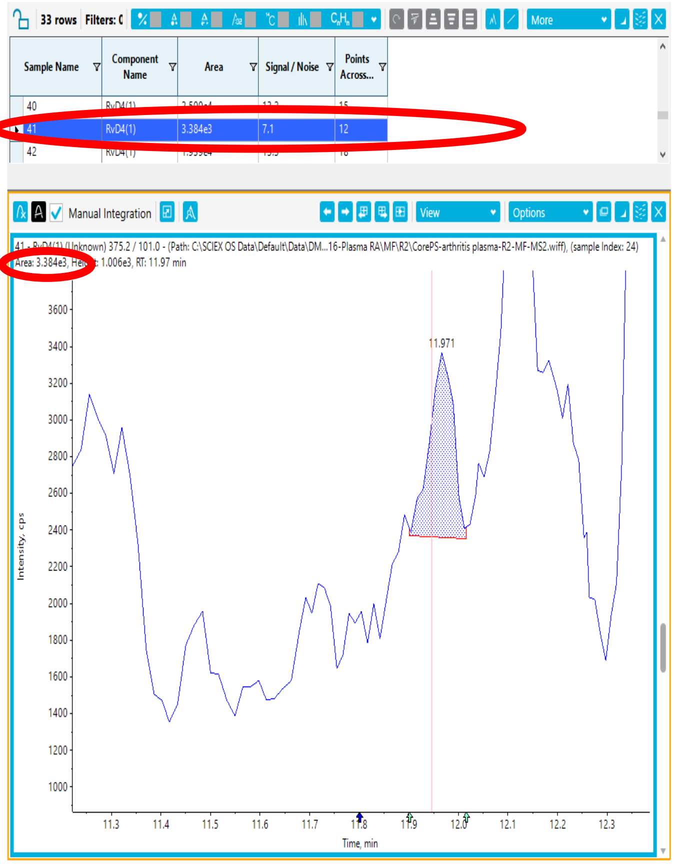


#### RvD5

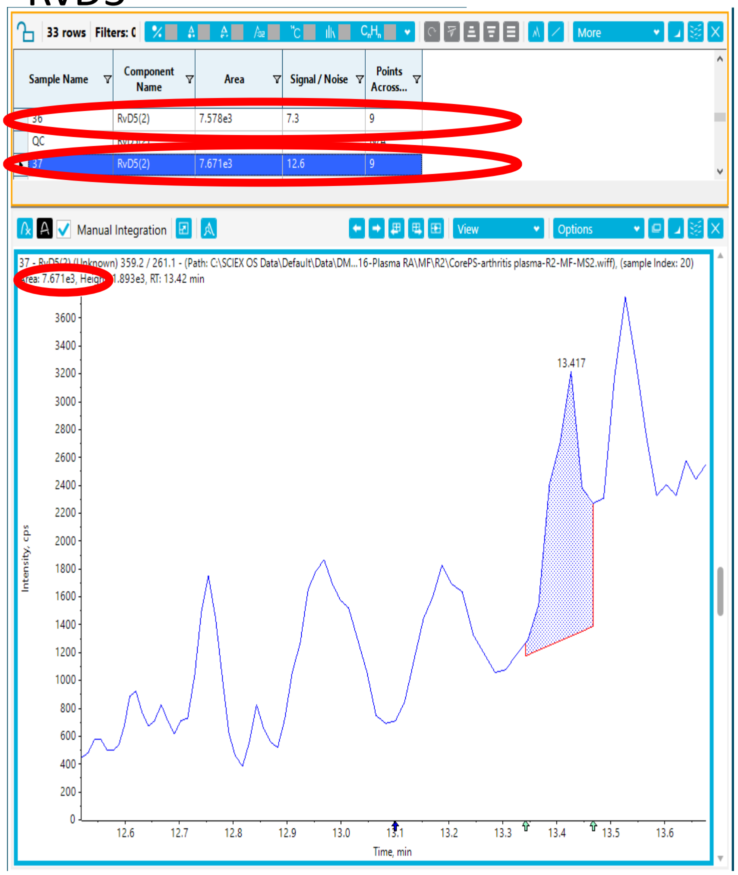

### Supplemental Figure 2

RvD6

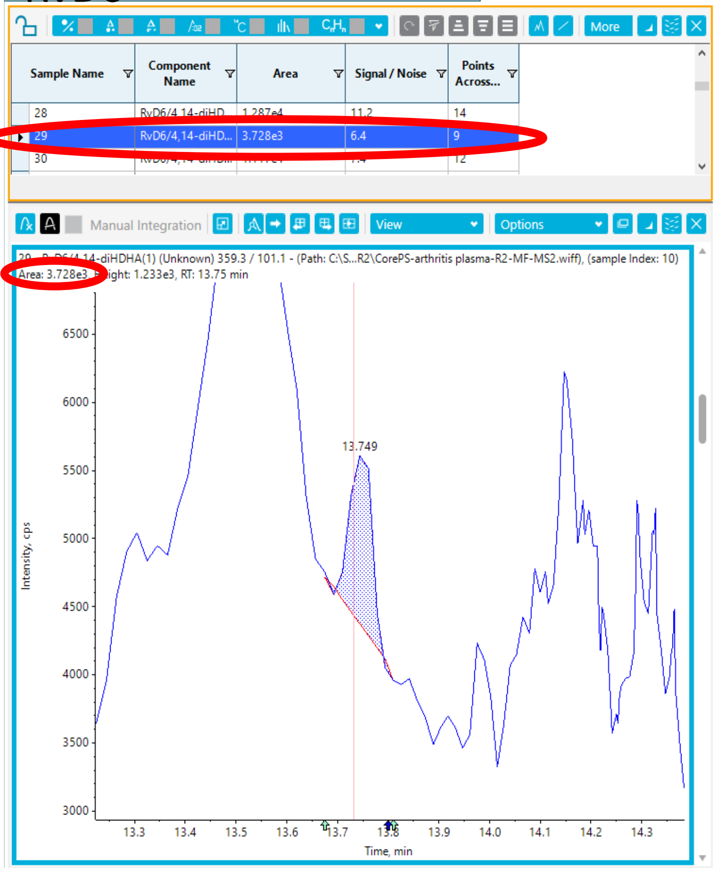


MaR1

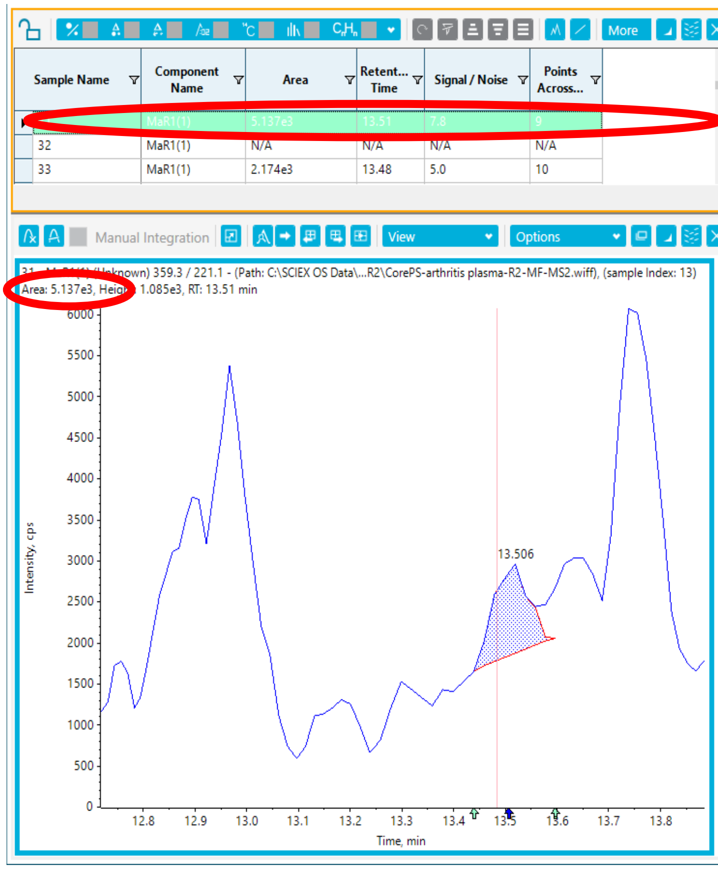


4S, 14S-diHDHA

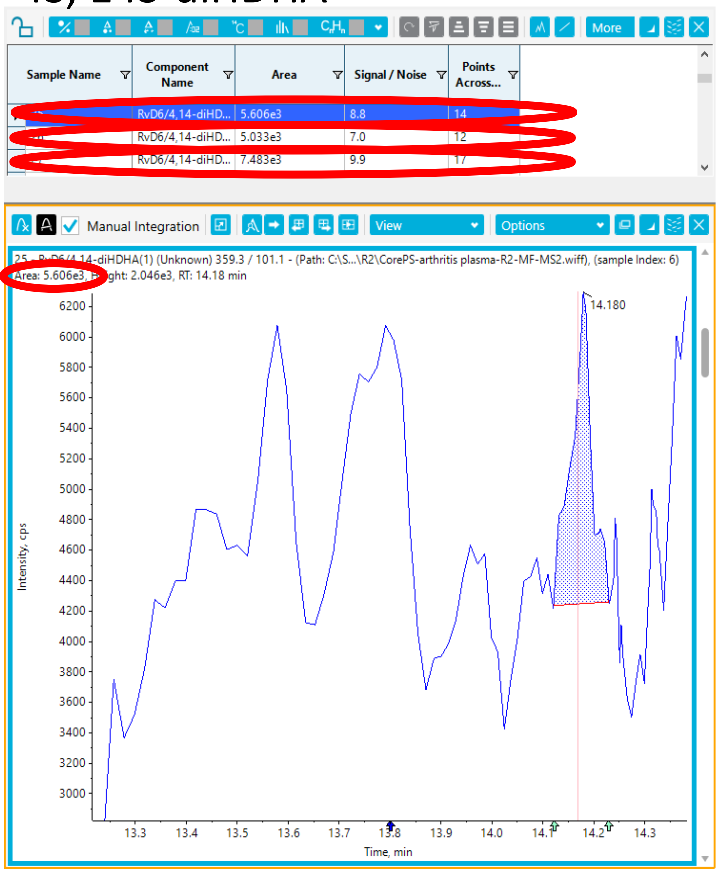


RvT1

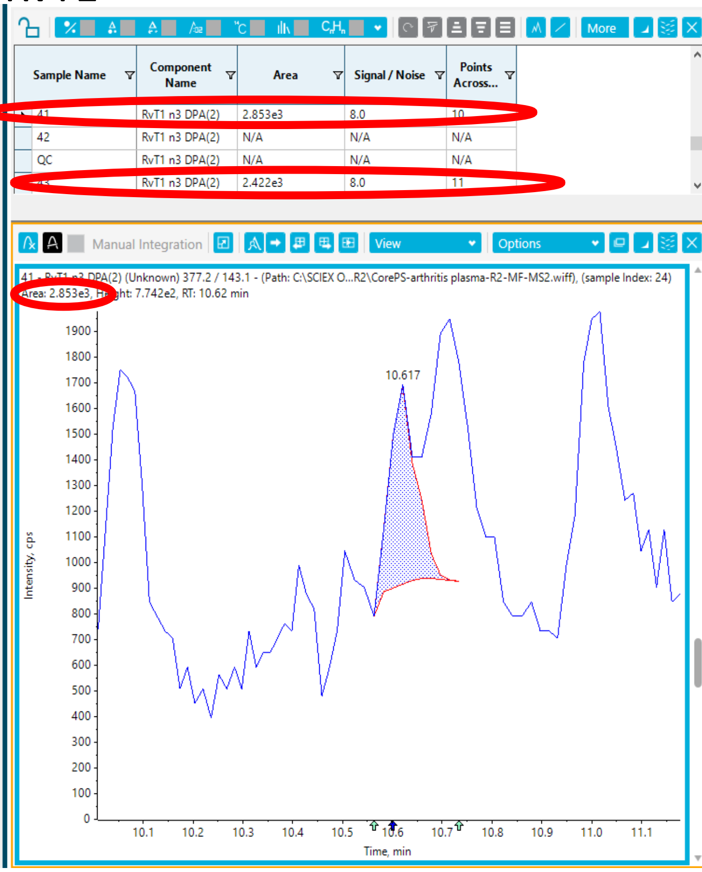

### Supplemental Figure 2

RvT4

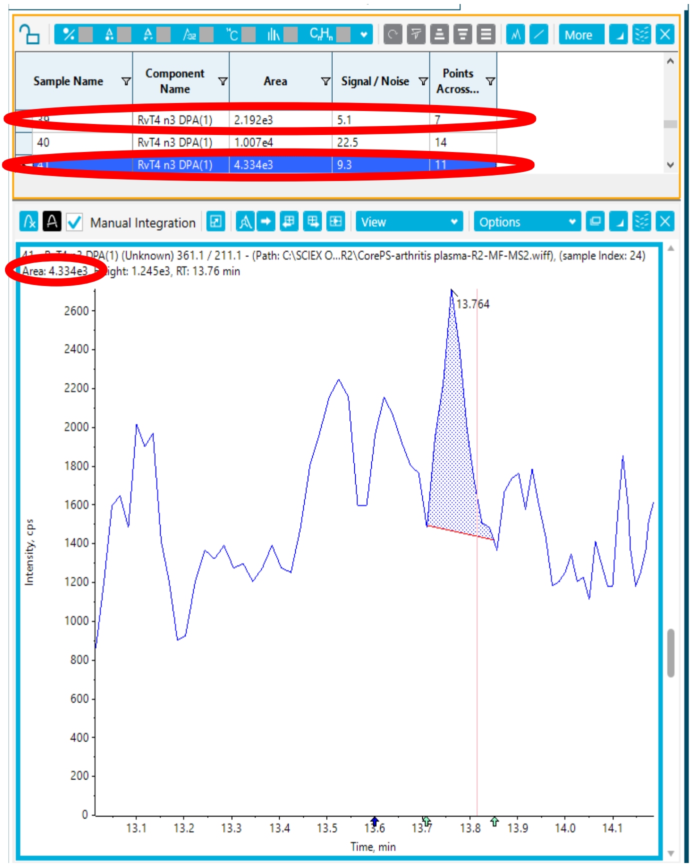


PD1n-3 DPA

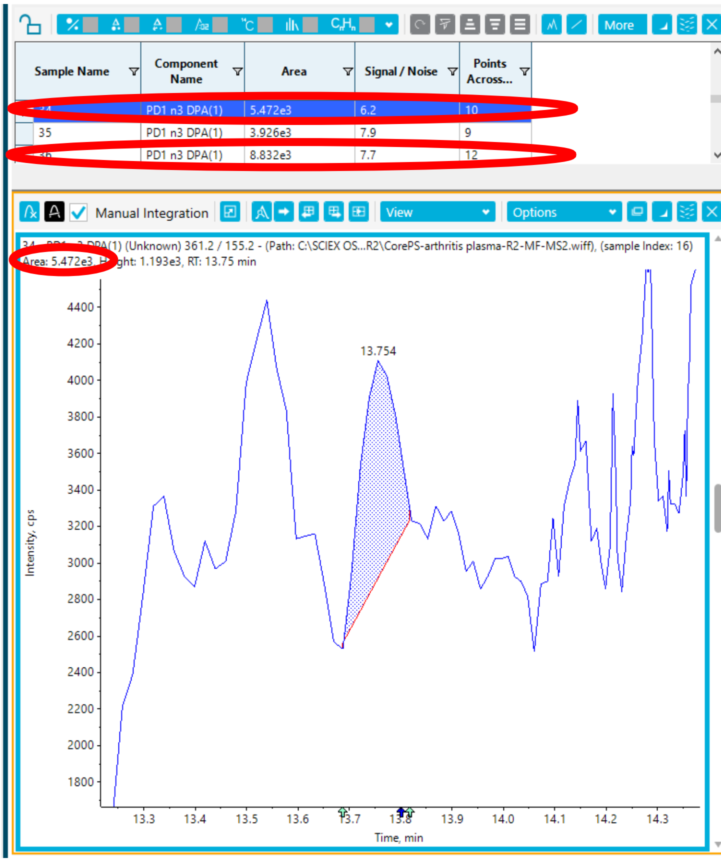


10S, 17S,diHDPA

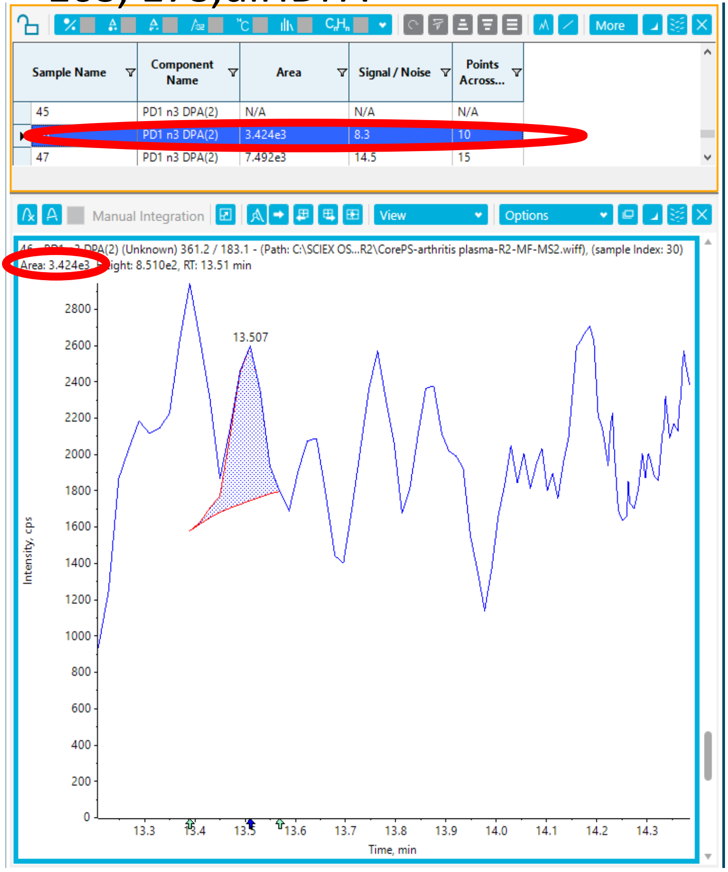


RvD1n-3 DPA

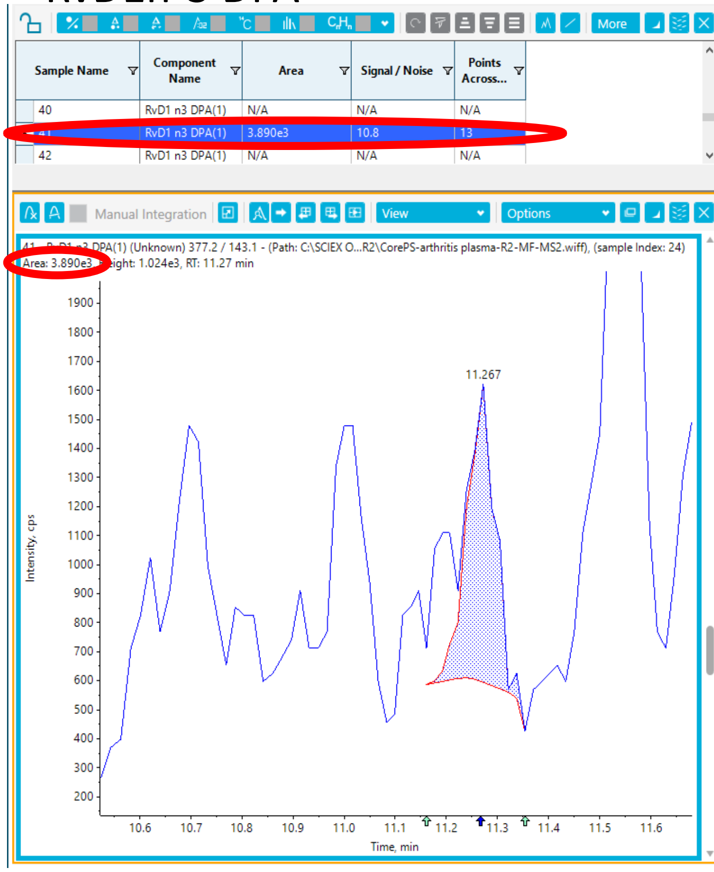

### Supplemental Figure 2

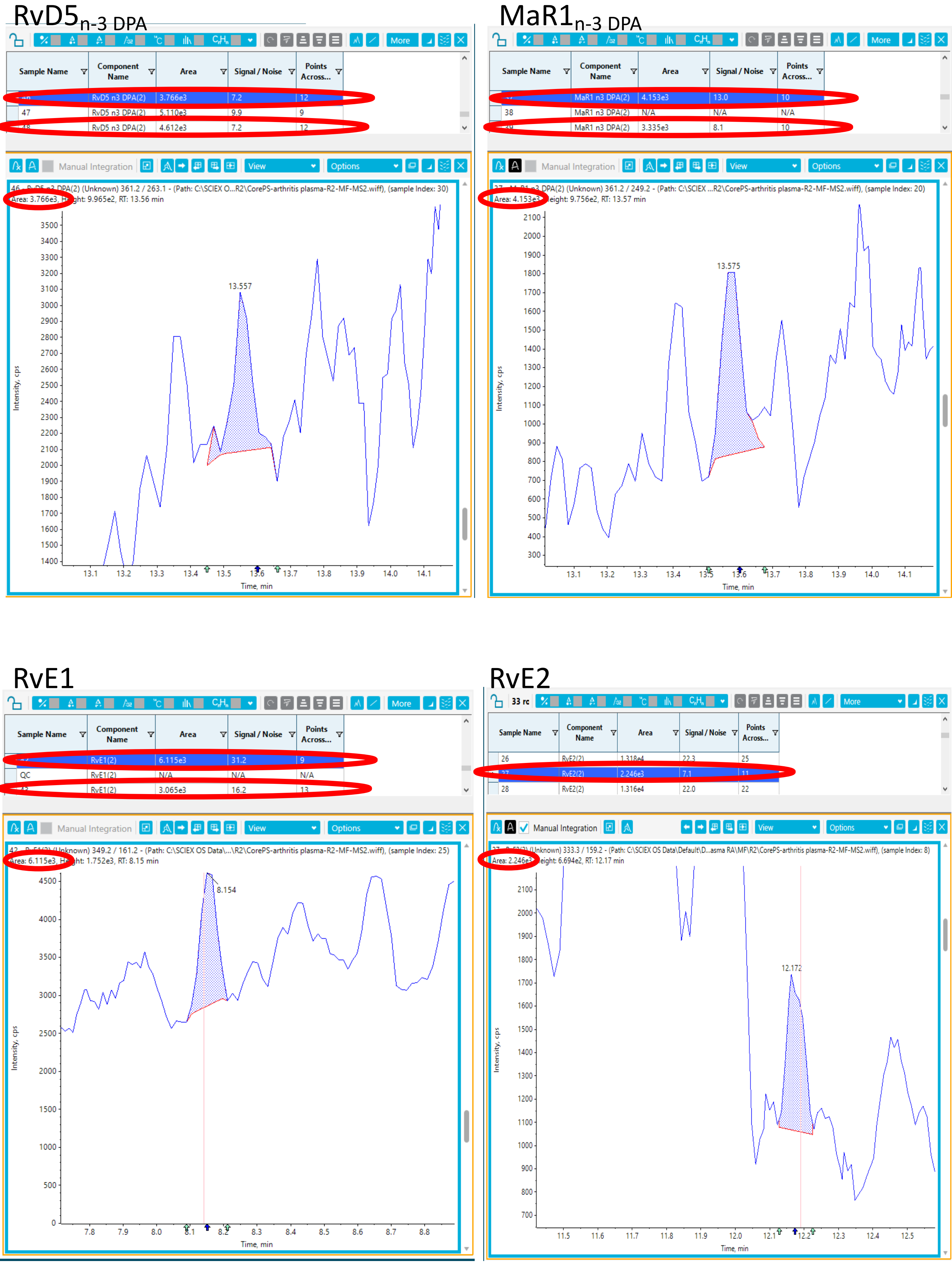

### Supplemental Figure 2

#### RvE3

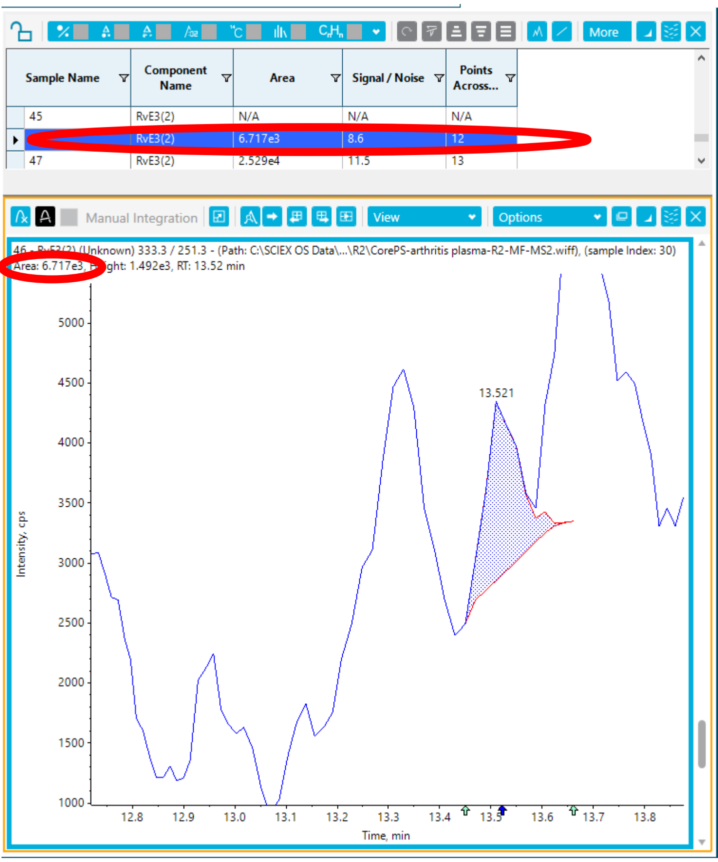


#### $LXB_4$

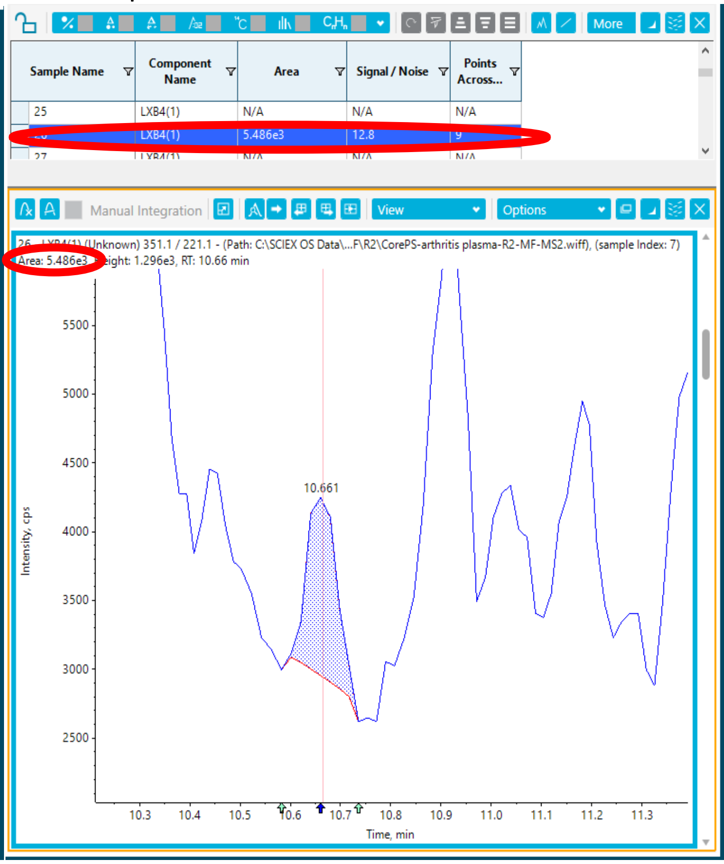


#### 10S, 17S-diHDPA

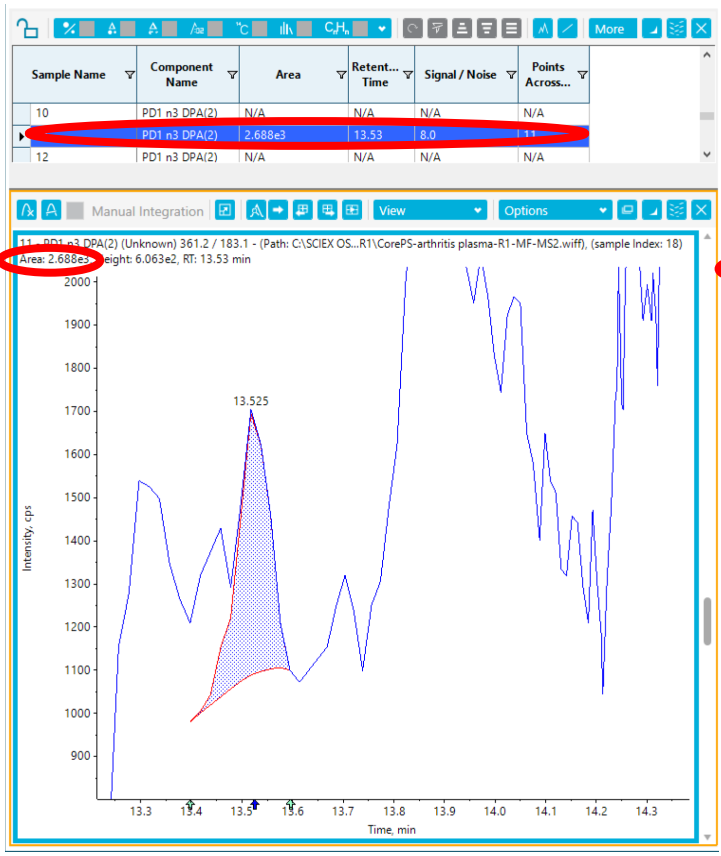


#### 14-oxo-MaR1

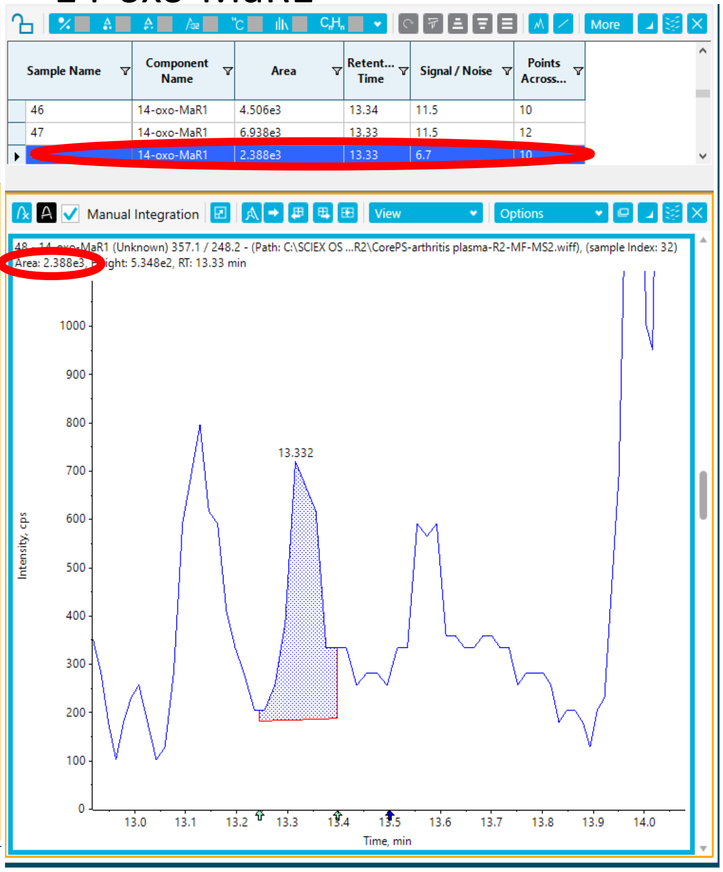

### Supplemental Figure 2

RvT2

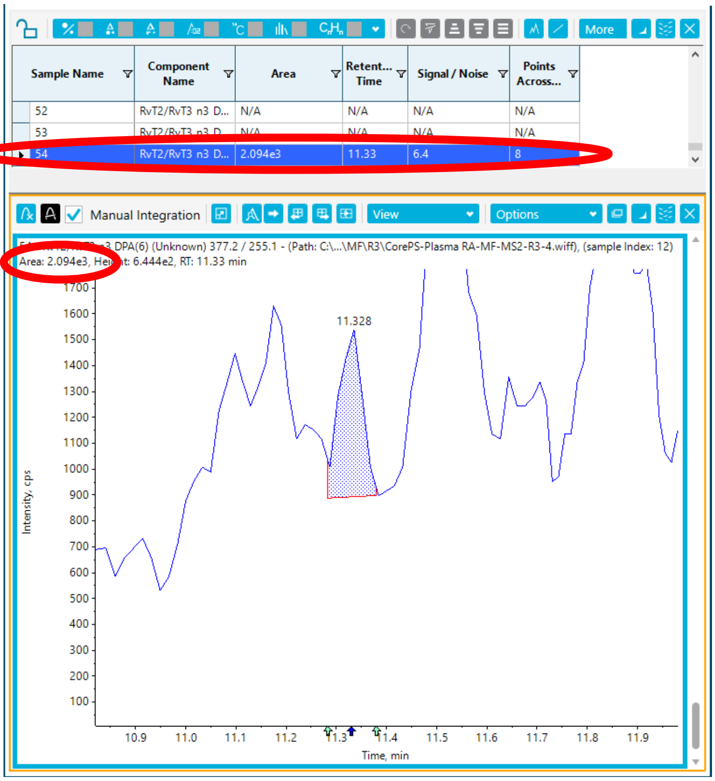


5S, 15S-diHETE

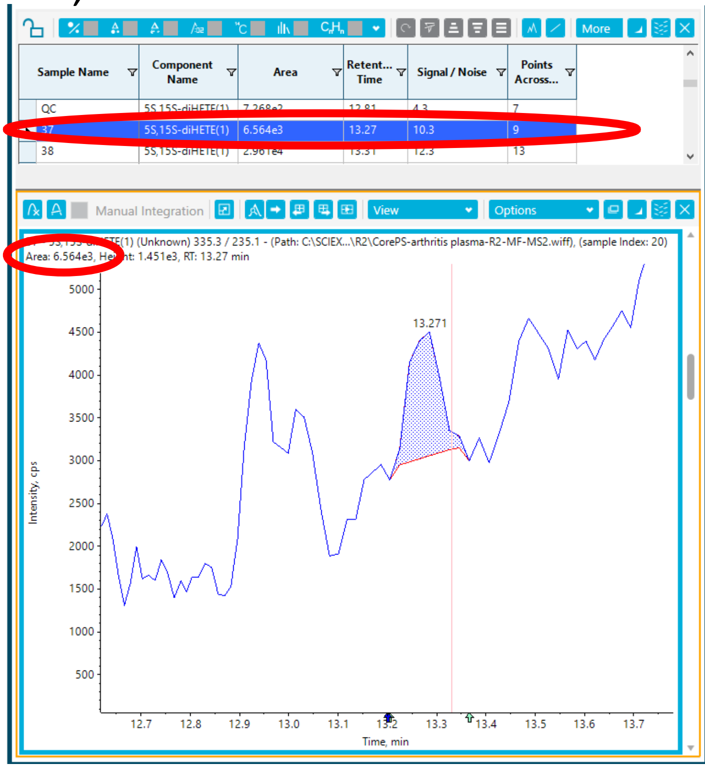

### Supplemental Figure 3

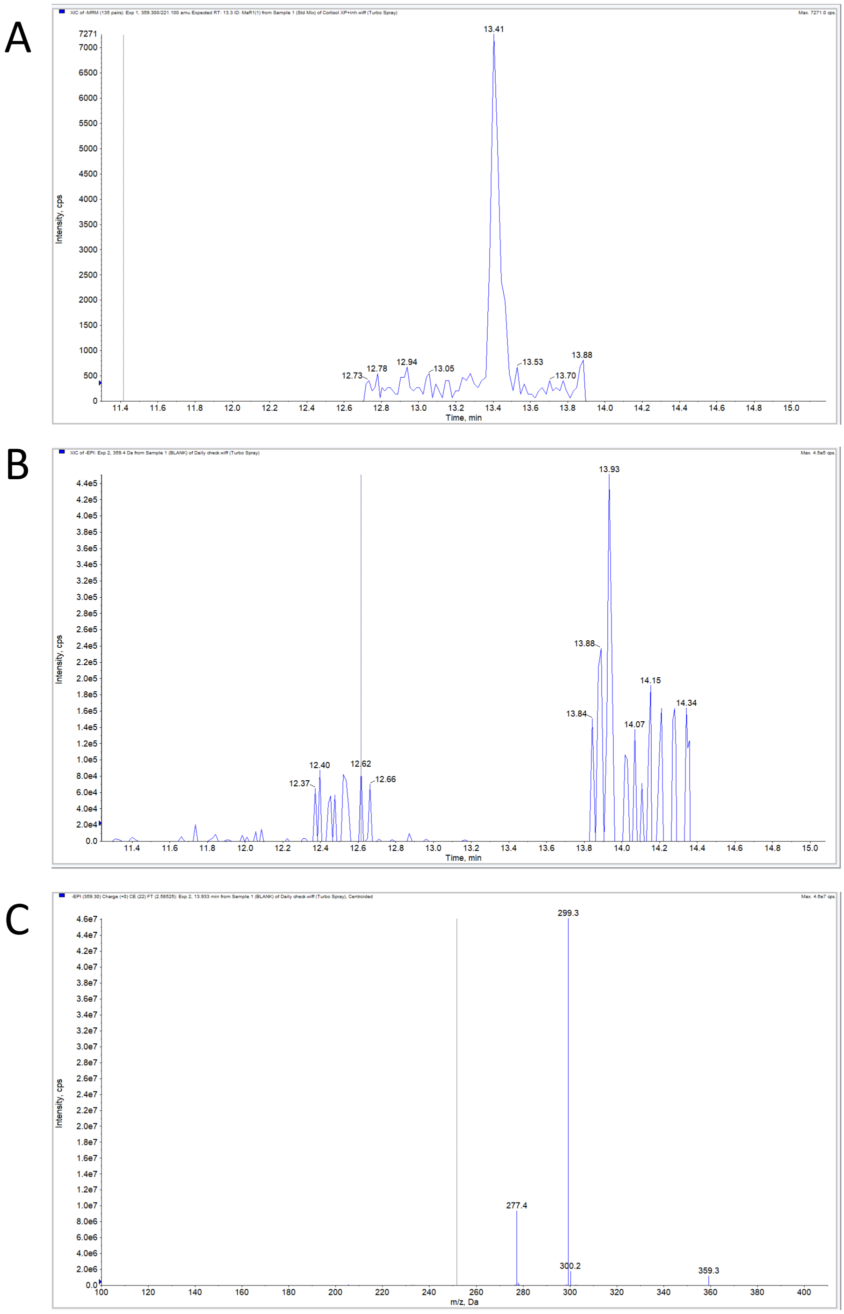

Supplemental Figure 4

RvD1

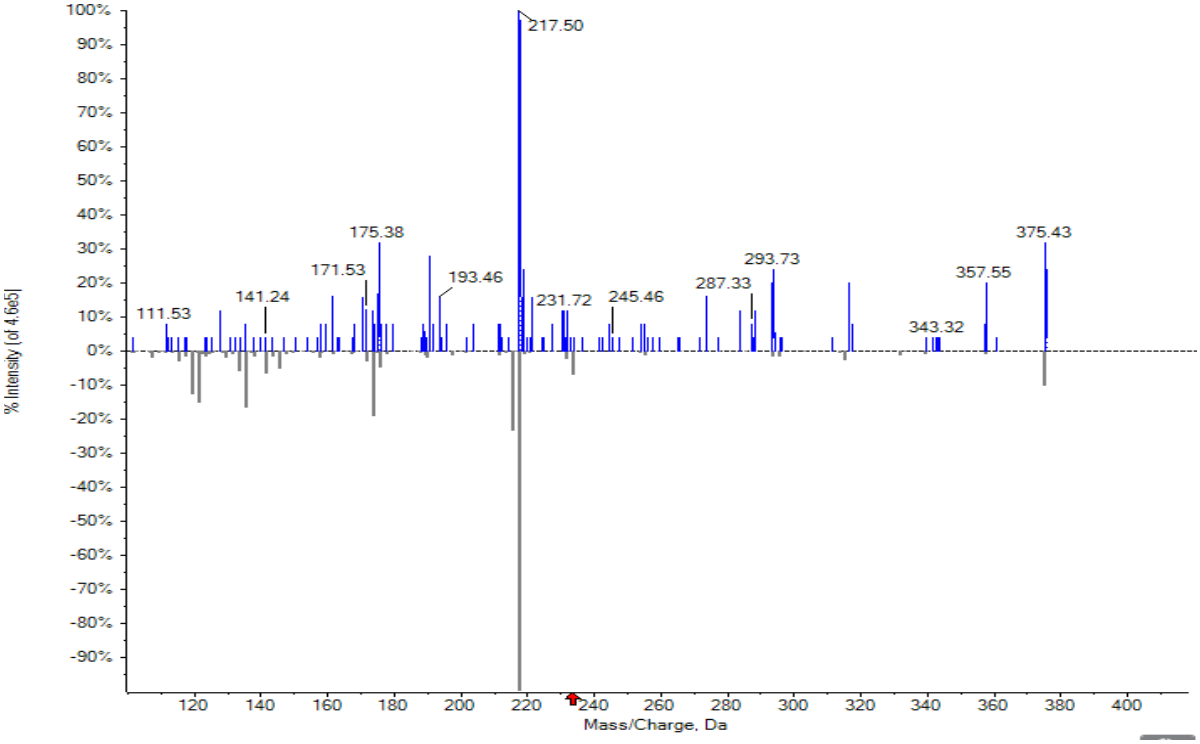


RvD4

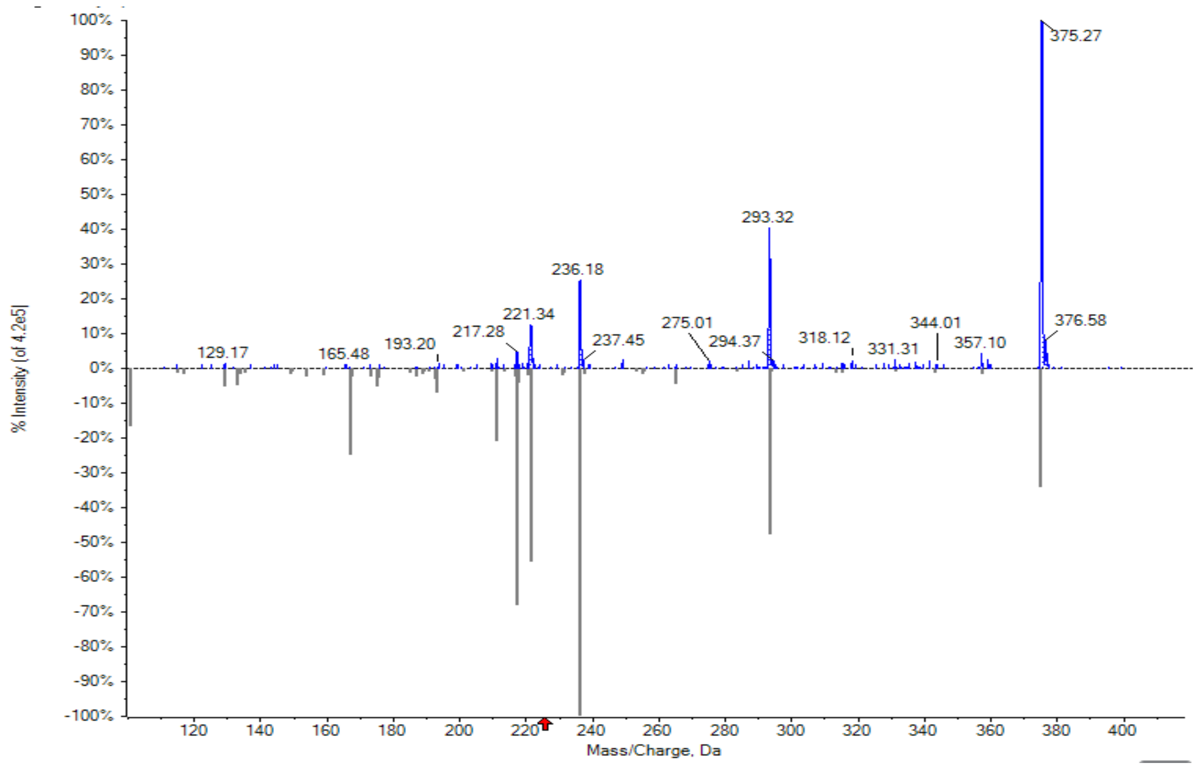

Supplemental Figure 4

RvD2

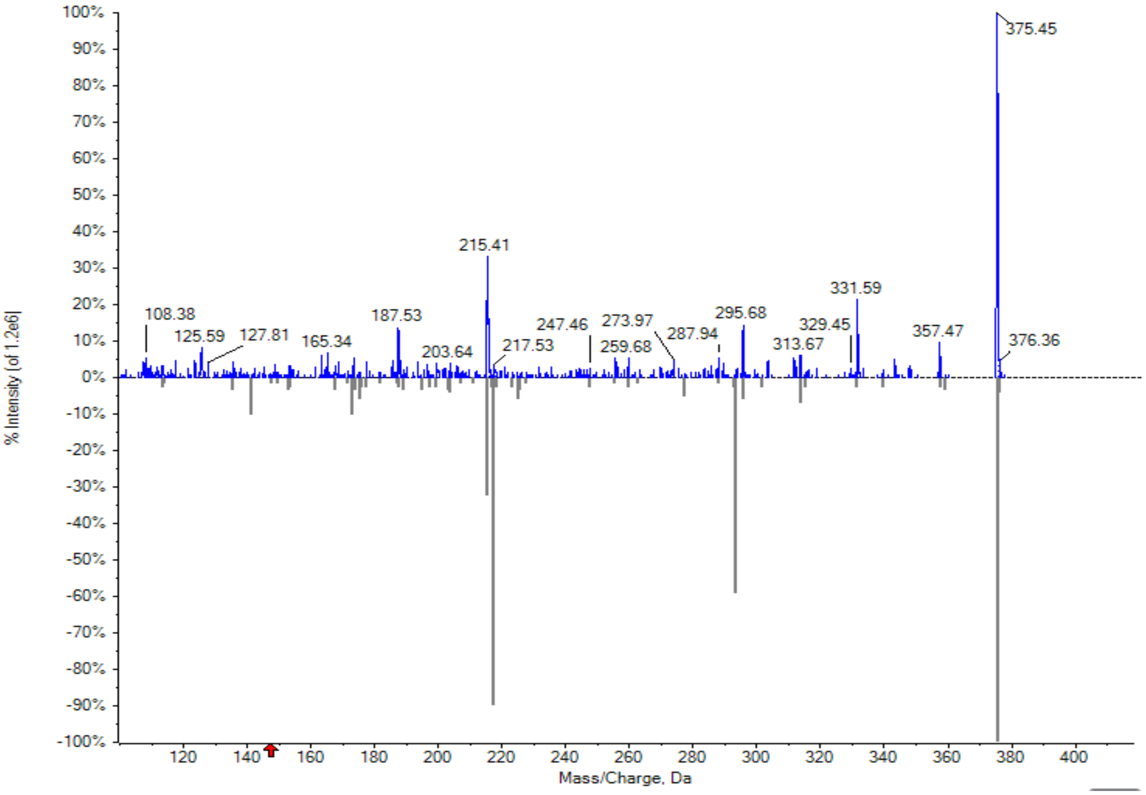


10S,17S-diHDPA

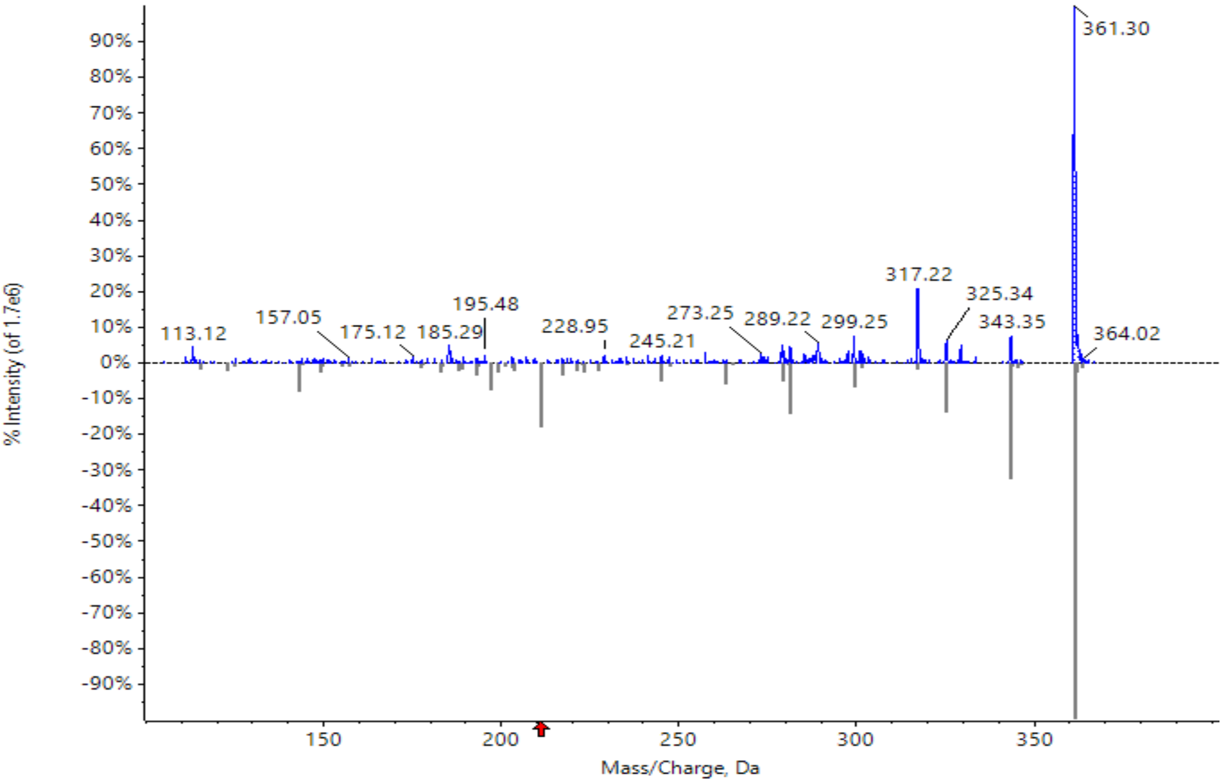

Supplemental Figure 4

RvD3

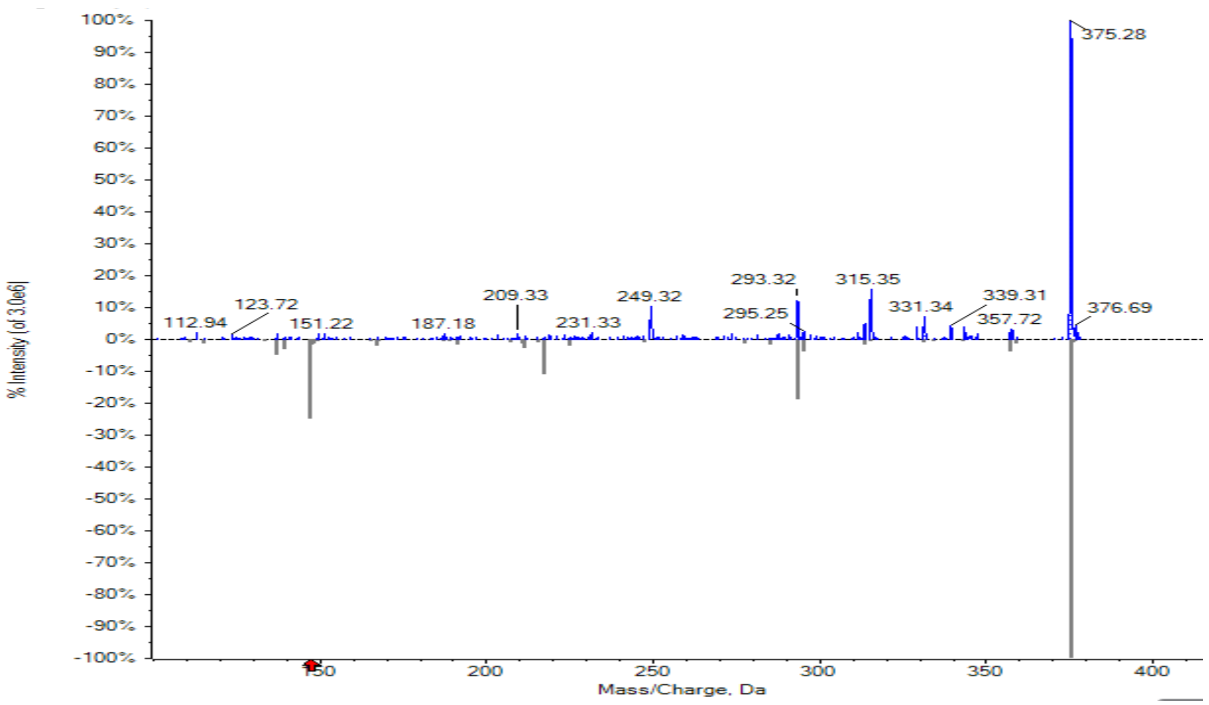


RvE3

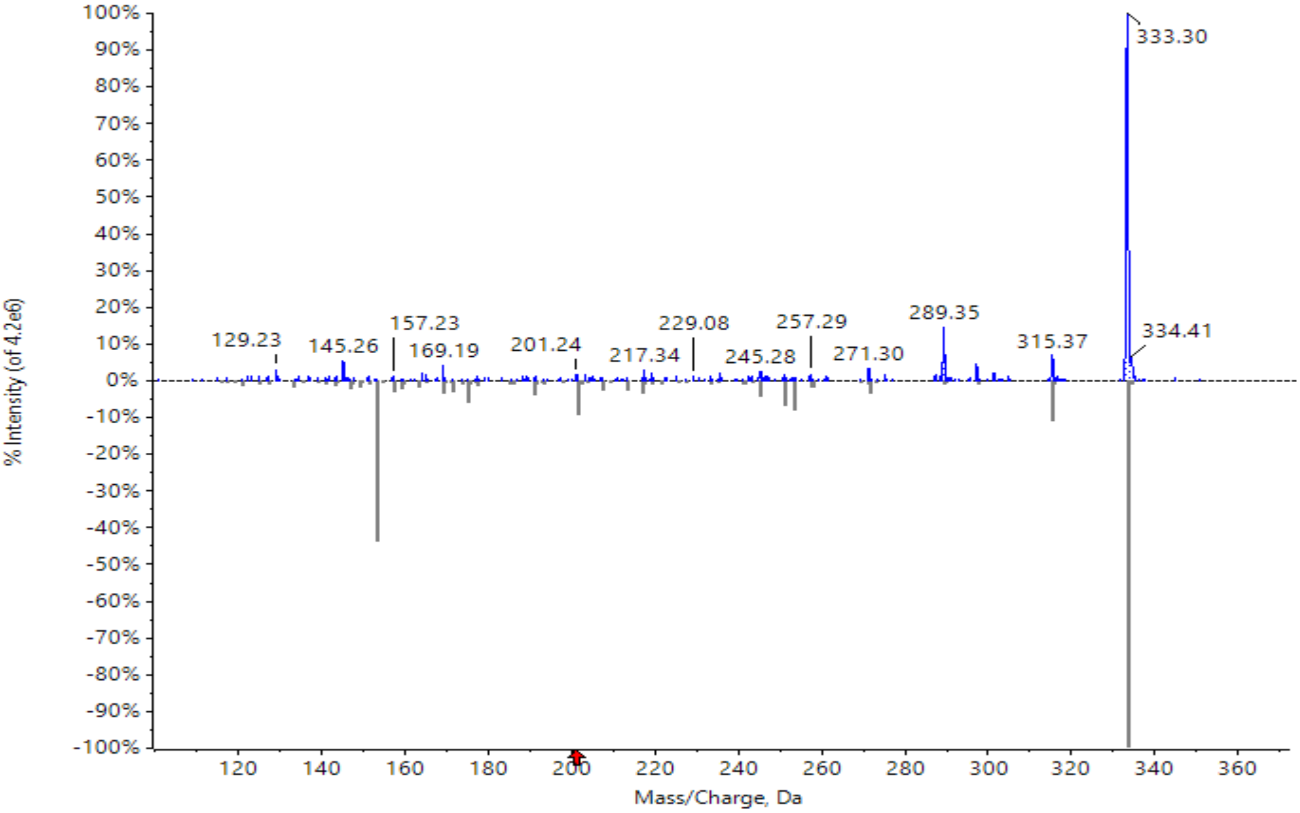

Supplemental Figure 4

$LXA_4$

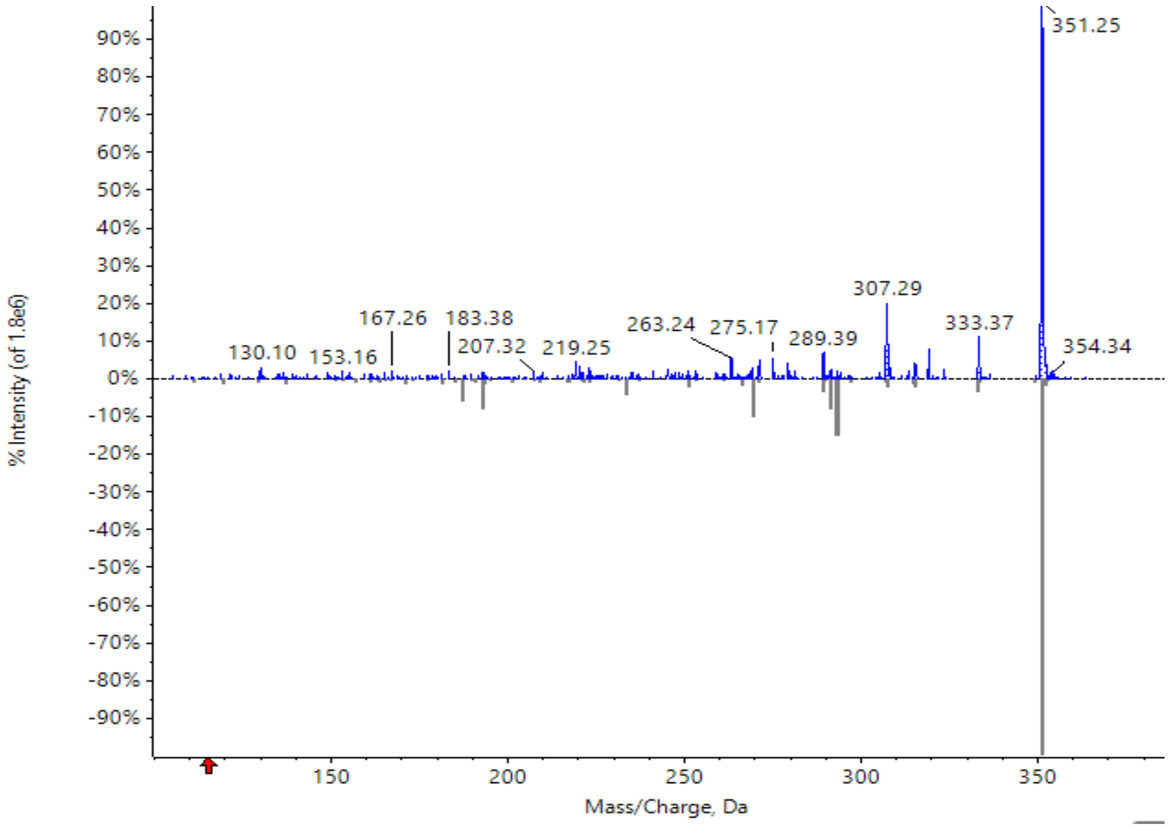


MaR1

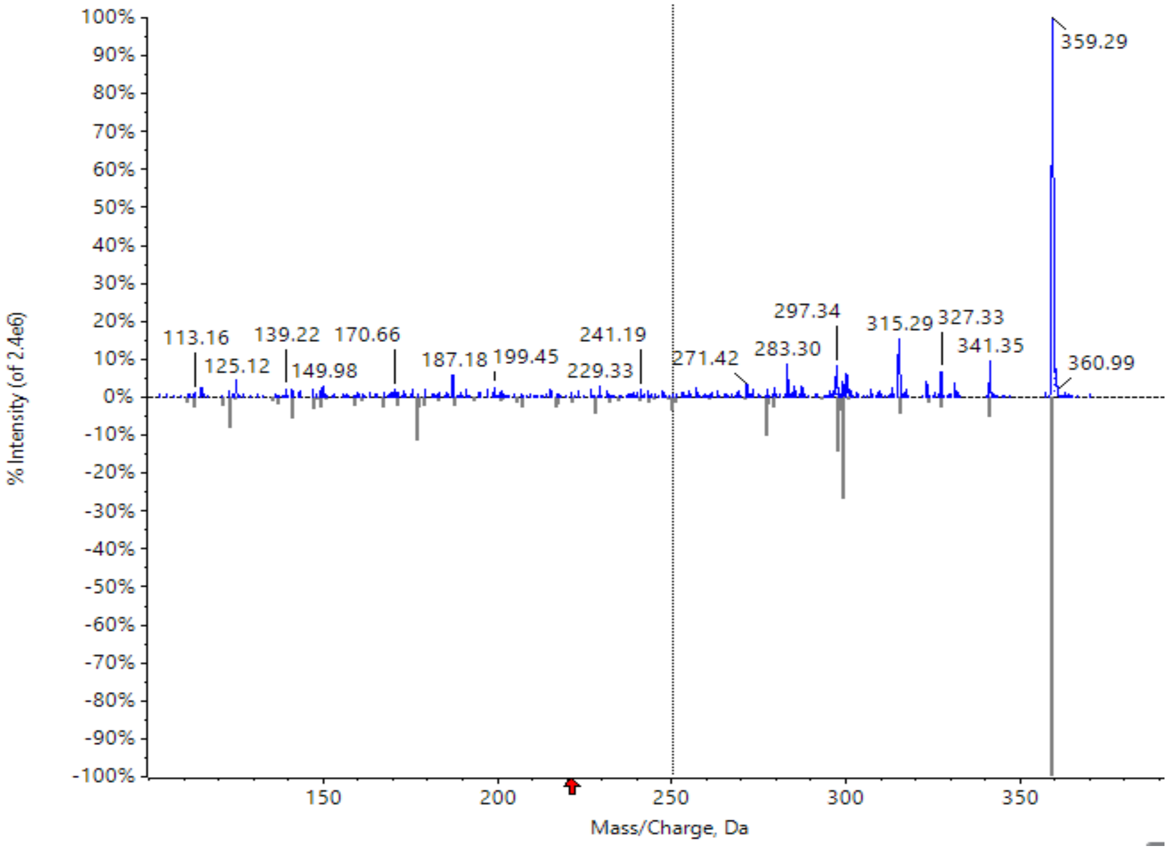
